# Early-life antibiotic exposure leads to gut microbial dysbiosis associated with neurodevelopment and neuroregeneration

**DOI:** 10.1101/2024.10.17.618899

**Authors:** Qing Zhao, Pianpian Fan, Qie Gu, Xiaoyu Mo, Xuemei Tan, Xinying Zhou, Fan Yang, Xiang Zhou, Qingchu Li

## Abstract

Early-life is a critical period for neurodevelopment and establishment of gut microbiota, which are highly susceptible to antibiotic exposure. Here, we aimed to investigate association between gut microbiota and metabolites with neurodevelopment and neuroregeneration in early-life after antibiotic exposure. To induce microbial dysbiosis in offspring, a broad-spectrum antibiotic cocktail was administered to mice dams. The prefrontal lobe, spinal cord tissues and gut sample of offspring were collected at different time with or without spinal cord crush injury. 16S rRNA sequencing and metabolomics were performed to analyze the gut microbiota and metabolites. NG2 glia were cultured and treated by mitochondrial fusion and fission compounds and metabolites. Maternal antibiotic exposure significantly affected neuronal maturation, NG2 glia proliferation, prefrontal cortex myelination and injured spinal cord neuroregeneration of offspring. Mice in antibiotic exposure group exhibited disruption of gut microbiota and metabolites, along with lower Ace, Chao and Shannon indexes, higher relative abundance of *Enterobacter* and *Escherichia_Shigella,* lower *lactobacillus* and *Streptococcus* genus, and downregulated Cyclo(Arg-Gly-Asp-D-Phe-Val), Ampicilloyl, DG(8:0/i-19:0/0:0), Petasinine and 6-Hydroxymelatonin glucuronide. These metabolites were enriched in alpha-Linolenic acid metabolism, phenylalanine, tyrosine and tryptophan biosynthesis, linoleic acid metabolism. 6-Hydroxymelatonin glucuronide, PC(20:3/0:0), PC(20:2/0:0), Ameltolide, DG(8:0/i-19:0/0:0), Inproquone, and Chorismate were the cores of networks between gut microbiota and metabolites and negatively associated with neurodevelopment and neuroregeneration. Additionally, gut microbiota metabolites promoted NG2 glia differentiation, partly through reversing the effects of mitochondrial fusion/fission compounds. Totally, antibiotic exposure in early-life changed the composition, abundance, and metabolites of gut microbiota in offspring, which was associated with neurodevelopment and neuroregeneration.

## Introduction

As one of the largest producers and consumers of antibiotics in the world, the consumption of antibiotics in China is increasing year by year. Studies showed that China consumed 162,000 tons of antibiotics in 2013, 48% of which were used for human beings and 52% for livestock and poultry (1). As a result, thousands of tons of antibiotics were released into the environment every year, resulting in serious antibiotic pollution (2). In our previous study, we found that urine samples collected from mothers during the third trimester of pregnancy and within 3 days after birth in an established cohort were extensively and severely exposed to antibiotics (3).

Antibiotics can be transported from mother to fetus and infant through the placenta and breast milk (4), leading to antibiotic exposure in early-life. Early-life is a critical period for neurodevelopment, especially during the fetal period and within 2 years after birth, which is highly susceptible to antibiotic exposure (5). Studies have shown that the use of antibiotics during pregnancy can increase the risk of attention deficit and hyperactivity disorder (ADHD) in children, leading to an increased risk of brain function damage in children and a significant reduction in overall cognition and language comprehension (5). The use of antibiotics in children is also associated with an increased risk of autism (ASD) and neurodevelopmental disorders (6). At the same time, the level of urinary antibiotics in early pregnancy is positively correlated with the increased risk of internalizing and externalizing problems in preschool children (7). A study based on 278 children aged 4-16 years showed that urinary levels of ciprofloxacin (classified as priority veterinary antibiotics, PVAs) were positively associated with increased risk of mental disorders, and increased levels of fluoroquinolones and priority veterinary antibiotics (PVAs) were positively associated with increased total difficulty scores (8). Animal studies have also found that maternal exposure to antibiotics during pregnancy may lead to social behavior disorder, anxiety and aggressive behavior in offspring rats, accompanied by adverse events such as hippocampal hypoplasia, limited neuronal proliferation, impaired synaptic plasticity and glial cell dysfunction (9). Our previous study also observed that the urine antibiotic level of infants was positively correlated with the increase of head circumference (3).

Early-life is also a critical window period for the establishment of gut microbiota, which is highly susceptible to the influence of antibiotics (10). Gut microbiota has been closely associated with neural development and brain function, and consequently, perturbation of the microbiota is implicated in several neurological disease (11, 12). Fetal life and infancy are crucial periods for individual neurodevelopment (13). The fact that the establishment and maturation of the infant gut microbiota co-occurs in parallel with key neurodevelopmental process (e.g., myelination, synaptogenesis) has led to realization that the early postnatal period represents a critical time-window for gut microbiota–brain interactions. Animal studies have indicated that even short-term changes in microbiota composition during the first few weeks after birth could have subtle but enduring effects on brain neurophysiology in early-life (14). While it is crucial to identify the association between the gut microbiota and neurodevelopment in the early-life.

Traumatic spinal cord injury (SCI) has various devastating consequences for the physical, financial, and psychosocial well-being. Normal spinal cord physiology involves interactions among different cell types such as astrocytes, neurons, and oligodendrocytes. After SCI, these multicellular interactions are interrupted and disorganized, leading to an impaired recovery (15). Neuroregeneration after SCI still remains a challenge due to the complicated inflammatory microenvironment and neuronal damage at the injury area (16). Clinical studies have identified gut microbiota dysbiosis in male patients with chronic traumatic complete SCI (17). Gut microbiota imbalance induced by oral broad-spectrum antibiotics aggravated mice neurological damage and myelopathy after SCI (18). In contrast, feeding SCI mice with a commercial probiotic rich in lactic acid-producing bacteria (VSL#3) or fecal microbiota transplantation (FMT) triggered neuroprotection with improving motor and gastrointestinal functions (18, 19). These results indicate that gut microbiota plays an important role in affecting neurological function and neuropathological recovery after SCI (18). Previous studies have showed that gut microbial dysbiosis after traumatic brain injury modulates the immune response and impairs neurogenesis (20). While few studies explored the effect of gut microbiota on neuroregeneration after SCI in early-life after antibiotic exposure.

Microbiota metabolites are compounds produced by intestinal microbial metabolism, which play an important role in neurodevelopment and neurodegenerative diseases (21). Recent studies have shown that trimethylamine oxide (TMAO), as a metabolite from gut microbiota in offspring rats, have shown to up-regulate the transcriptional expression of mitochondrial fusion-related genes(22). The developing brain is the most energy-intensive organ and relies heavily on mitochondria for energy metabolism. Therefore, mitochondria play a key role in neuronal development, neurotransmission and neuroplasticity (23, 24), and participate in the occurrence of a variety of nervous system diseases such as autism spectrum disorder (ASD) (25). Normally, mitochondria are in a highly dynamic state of change, often undergoing fission and fusion to maintain their structural and functional homeostasis (26), which in turn affects axon growth, synapse formation, and even neuronal apoptosis (27). Accordingly, mitochondrial fission/fusion homeostasis is a key factor affecting neurodevelopment. We hypothesized that gut microbiota metabolites might modulate the effects of early-life antibiotic exposure on neurodevelopment by perturb mitochondrial fission/fusion homeostasis.

Shifts in the gut microbiota led to altered metabolomic profiles, impacting the availability and diversity of nutrients and metabolites that may interact with the central nervous system. In the present study, we aimed to characterize the impact of antibiotic exposure in early-life induced gut microbiota disruption and metabolites on neurodevelopment in prefrontal lobe and neuroregeneration after SCI. We also explored whether the exposure of gut microbiota metabolites and mitochondrial fusion/fission compounds would affect the differentiation of NG2 glia.

## 2. Materials and methods

### 2.1 Animal modeling, grouping, and intervention

All procedures were carried out in accordance with protocols approved by the Animal Care and Use Committee of Southern Medical University (Guangzhou, China). The experimental procedures were designed to minimize the number of animals used as well as animal suffering. C57/BL6 mice (12 weeks of age) were obtained from Guangdong Zhiyuan Biotechnology Co., Ltd. Mice were housed on a 12/12-h light/dark cycle, and food and water were available ad libitum. The room temperature and humidity were appropriate. Mice were individually housed to prevent cross-colonization of mice in the same cage as previously described (18). Their offspring were randomly divided into the following 6 groups: 1) Control (CON) group (n=15), dams with no treatment. 2) antibiotics (ABX) group (n=15), dams were fed with antibiotics. 3) sham group (n=3), dams without antibiotics treatment, offspring were conducted with laminectomy but not spinal cord crush on 7^th^ day after birth. 4) sham+ABX group (n=3), dams were treated with antibiotics, offspring were conducted with laminectomy but not spinal cord crush on 7^th^ day after birth. 5) SCI group (n=12), dams without antibiotics treatment, while offspring were conducted with SCI on 7^th^ day after birth. 6) SCI and antibiotics (SCI+ABX) group (n=12), dams were fed with antibiotics, offspring were conducted with SCI on 7^th^ day after birth. The procedure for dams gavaged with antibiotics was described in the “Antibiotics administration” section below. Three or more independent experiments were performed.

### 2.2 Antibiotics administration

Dams (pregnant mice, 12 weeks of age) exposed to antibiotics as a proxy for evaluating the contribution of gut microbiota disturbance to early neurodevelopment and neuroregeneration. Antibiotics administration was described in the experimental flow chart (**Supplementary Figure 1)**. To induce microbial dysbiosis in offspring, a broad-spectrum antibiotic cocktail (ampicillin, neomycin, metronidazole and vancomycin) was administered to mice dams (C57/BL6, 12 weeks of age) from 1 week before until 1 week after giving birth. Dams in ABX, sham+ABX and SCI+ABX groups received oral broad-spectrum antibiotic cocktail as previously reported (19), including 0.2 g/L ampicillin (Cat# A9518, Sigma-Aldrich), 0.2 g/L neomycin (Cat# N6386, Sigma-Aldrich), 0.2 g/L metronidazole (Cat# PHR1052, Sigma-Aldrich), and 0.1 g/L vancomycin (Cat# V2002, Sigma-Aldrich) in saline twice daily for 2 weeks, which sustained from 1 week before giving birth until 1 week after giving birth. Control groups of sham and SCI received the same amount saline for the same duration as the groups mentioned above. Dams resumed regular drinking water for the remainder of experiment. Antibiotics were renewed every two days.

### 2.3 Spinal cord crush model

The crush model was performed as we previously reported (28). Briefly, juvenile mice were deeply anaesthetized with isoflurane evaporated in a gas mixture containing 70% N_2_O and 30% O_2_ through a nose mask (28). The back skin was shaved and cleaned with 75% alcohol. A laminectomy at thoracic 10 was performed to remove the part of the vertebra overlying the spinal cord, exposing a circle of dura through an operating microscope. The crush model was performed at thoracic with No. 5 Dumont forceps fixed on the stereotaxic apparatus (RWD Life Science Co.,Ltd, Shenzhen, China) and persisted for 3 seconds, and the wound was sutured (28). The paralysis of both hindlimbs occurred after awakening were indicators of a successful crush model. Then the juvenile mice were returned to the dams’ cage for feeding after crush during the experiment (28).

### 2.4 Immunofluorescence staining and analysis

Immunofluorescence staining procedures were conducted as we previously described (28). For mice offspring in control and ABX group, the prefrontal lobes of brain were collected on the 1^st^, 3^rd^, 7^th^, 14^th^, 21^th^ days after birth (**Supplementary Figure 1)**. For mice offspring in sham and sham+ABX group, the thoracic spinal cord was collected on the 14 days post injury (dpi) after laminectomy. For mice offspring in SCI and SCI+ABX group, the lesion area of spinal cord was collected on the 1dpi, 3dpi, 7dpi, and 14dpi after SCI. After post-fixation and cryoprotection, a dissected 6 mm segment of spinal cord centred around the injury epicentre, and the prefrontal lobe of brain, was coronally sectioned at 12 µm thickness and thaw-mounted onto Superfrost Plus slides (Citotest Labware Manufacturing Co., Ltd.) (28). The primary antibodies used were as follows: TUJ1 (Cat# MAB1637, Millipore, 1:500), Myelin basic protein (MBP, Cat# MAB386, Millipore, 1:500), MBP (Cat# 78896T, CST, 1:500), Neural/glial antigen 2 (NG2, Cat# AB5320, Millipore, 1:500), PDGF Receptor alpha/PDGFRA (Cat# sc-398206, Santa Cruz Biotechnology, 1:100), MAP2 (Cat# 8707T, CST, 1:200); the secondary antibodies were Alexa® Fluor 488 (Cat#A-32814, Invitrogen, 1:500), Alexa® Fluor 594 (Cat# ab150176, Abcam, 1:500), and Alexa® Fluor 647 (Cat# A-32795, Invitrogen, 1:500). The nuclei were stained with 4′,6-diamidino-2-phenylindole (DAPI), and fluorescence images were taken and assembled as we previously reported (28).

For immunofluorescence analysis of prefrontal lobe and spinal cord, three sections were randomly selected, then the 400 μm × 400 μm images (prefrontal lobe > 45 images per groups, spinal cord > 15 images per groups) were exported as single channel images (28). Image-J software with customized macros was used to quantify the fluorescence intensity of proteins in the images mentioned above. All images of prefrontal lobe and spinal cord sections were modified through the same procedures. 3 mice per group were tested.

### 2.5 Gut sample collection

For mice offspring in control and ABX group, the gut sample were collected on the 1^st^, 3^rd^, 7^th^, 14^th^ days after birth (**Supplementary Figure 1)**. For mice offspring in sham and sham+ABX group, the gut sample were collected on the 14dpi after sham treatment. For mice offspring in SCI and SCI+ABX group, the gut sample were collected on the 1dpi, 3dpi, 7dpi, and 14dpi after SCI. Briefly, the fur of mice was removed, and then the surface of each mouse was sterilized using 75% ethanol and then the intestine from the duodenum to the colon with feces were gently removed with sterile tweezer and surgical blade, and moved into a sterile DNAase free tube and stored at −80°C until processing.

### 2.6 DNA Extraction, 16S rRNA gene sequencing and analysis

16S rRNA gene sequencing of the gut samples was carried out as previously reported (29). Briefly, the samples were subjected to polymerase chain reaction (PCR) amplification, after which the amplicons were purified and subjected to paired-end sequencing and the raw data was evaluated. Information on sequencing analysis is provided in the **Appendix**.

### 2.7 Examination and analysis of untargeted metabolomics

Due to limited samples collected, gut samples in SCI mice on 1dpi, 3dpi, 7dpi and 14dpi, as well as SCI+ABX mice on 7dpi and 14dpi were analyzed for metabolomics. The detailed protocol for fecal sample preparation for LC-MS analysis is provided in the appendix. Chromatographic separation of the metabolites was performed on a Thermo UHPLC system equipped with an ACQUITY BEH C18 column (100 mm × 2.1 mm i.d., 1.7 µm; Waters, Milford, USA). The details were supported in **Appendix**. Partial least-squares discriminant analysis (PLS-DA) and orthogonal partial least squares discriminate analysis (OPLS-DA) supervised pattern recognition methods were used for statistical analysis to determine global metabolic changes between comparable groups. Variable importance in projection (VIP) ranks the overall contribution of each variable to the OPLS-DA model. Variables with VIP > 1.0 and p < 0.05 were considered relevant for group discrimination. Finally, the differential metabolites were annotated through the metabolic pathways in the KEGG database (https://www.kegg.jp/kegg/pathway.html) to obtain the pathways involved in the differential metabolites. Further information is provided in the **Appendix**.

### 2.8 Gut microbiota–metabolite correlation analysis

We firstly identified the differential metabolites in the SCI and SCI+ABX groups, which are engaged in important metabolic pathways, via correlation analysis. Then, we identified the top 30 most abundant species in the gut microbiota in the groups. Lastly, we used Spearman correlation analysis and heatmaps to assess the correlation between gut microbiota and metabolite content.

### 2.9 NG2 glia culture and treatment

We treated NG2 glia, also named oligodendrocyte precursor cells (OPCs), with metabolites and mitochondrial fusion/fission compounds. OLN-93 cells (rat NG2 glia, Cat # CL-030r) and MO3.13 cells (Human NG2 glia, Cat # CL-526h) were obtained Wuhan Saios Biotechnology Co., LTD and were cultured in the NG2 glia proliferation medium as we previously reported (28). In brief, OLN-93 cells and MO3.13 cells were cultured in a 100 mm non-coated Corning dish with NG2 glia proliferation medium (neurobasal medium (Cat# 10888022, Gibco), 10 ng/ml PDGF-AA (Cat# 315-17, Peprotech), 10 ng/ml bFGF (Cat# 45033, Peprotech), 2 mM glutamine (Cat# G7513, Sigma), 1% penicillin/streptomycin, 5 ng/ml insulin (Cat# P3376-100 IU, Beyotime Institute of Biotechnology, China), 20 ug/ml NT3 (Cat# 450-03-10, Peprotech), and 2% B27 supplement (Cat# A3582801, Gibco)) (28).

After reaching 70–80% confluency at 3 days, NG2 glia (OLN-93 and MO3.13 cells) were treated by mitochondrial fusion and fission compounds and metabolites for 7 days, including mitochondrial division inhibitor Midivi-1 (Cat# HY-15886, MCE, 10μM(26)), mitochondrial fusion promoter M1 (Cat# HY-111475, MCE, 5μM(26)) and metabolites (trimethylamine-N-oxide (TMAO, Cat# C3481, APExBIO, 1μM (11)), imidazole propionate (IP, Cat# 33458, Cayman, 1μM (11)), hippurate (HP, Cat# M1191, APExBIO, 1μm (11)). Cells were randomly divided into 6 groups: Control, DMSO, M1, Midivi-1, Metabolites (TMAO+IP+HP), Metabolites+M1, Metabolites+ Midivi-1 groups. After treatment for 7 days, NG2 glia were collected for immunocytochemistry. Images were obtained by Olympus fluorescence microscope. Three independent experiments were performed.

### 2.10 Statistical analysis

All experiments were conducted with three duplicates. All continuous data were shown as mean ± SEM. Data were analyzed using Student’s *t*-test for comparisons between two groups. Spearman index was used to evaluate the association between gut microbiota and markers for neurodevelopment or neuroregeneration. Data for littermates from the same group were averaged and presented in the main figures. P values < 0.05 were considered statistically significant. Data analyses were conducted using the Statistical Analysis System (SAS), version 9.4 (SAS Institute, Inc, Cary, NC, USA). Plots were generated using GraphPad Prism 8 software (GraphPad Software, San Diego, CA).

## 3. Results

### 3.1 Maternal exposure to antibiotics impacted on the brain neurodevelopment and gut microbiota of offspring

Immunofluorescence staining was used to demonstrate the impact of maternal exposure to antibiotics on the neurodevelopment in prefrontal lobe (**Figure 1**). Compared with CON group, we found that the offspring mice in ABX group had significant higher expression of neuronal marker (TUJ1) on the 1^st^, 7^th^, 14^th^, 21^st^ days after birth (**Figure 1A-B**), along with higher oligodendrocytes maturation and myelin formation (MBP), and NG2 glia proliferation (NG2) (**Figure 1A, 1C-D**). These results indicated that dams exposed to antibiotics would impact on the neuronal development and oligodendrocytes maturation in prefrontal lobe of their offspring mice.

**Figure 1.**
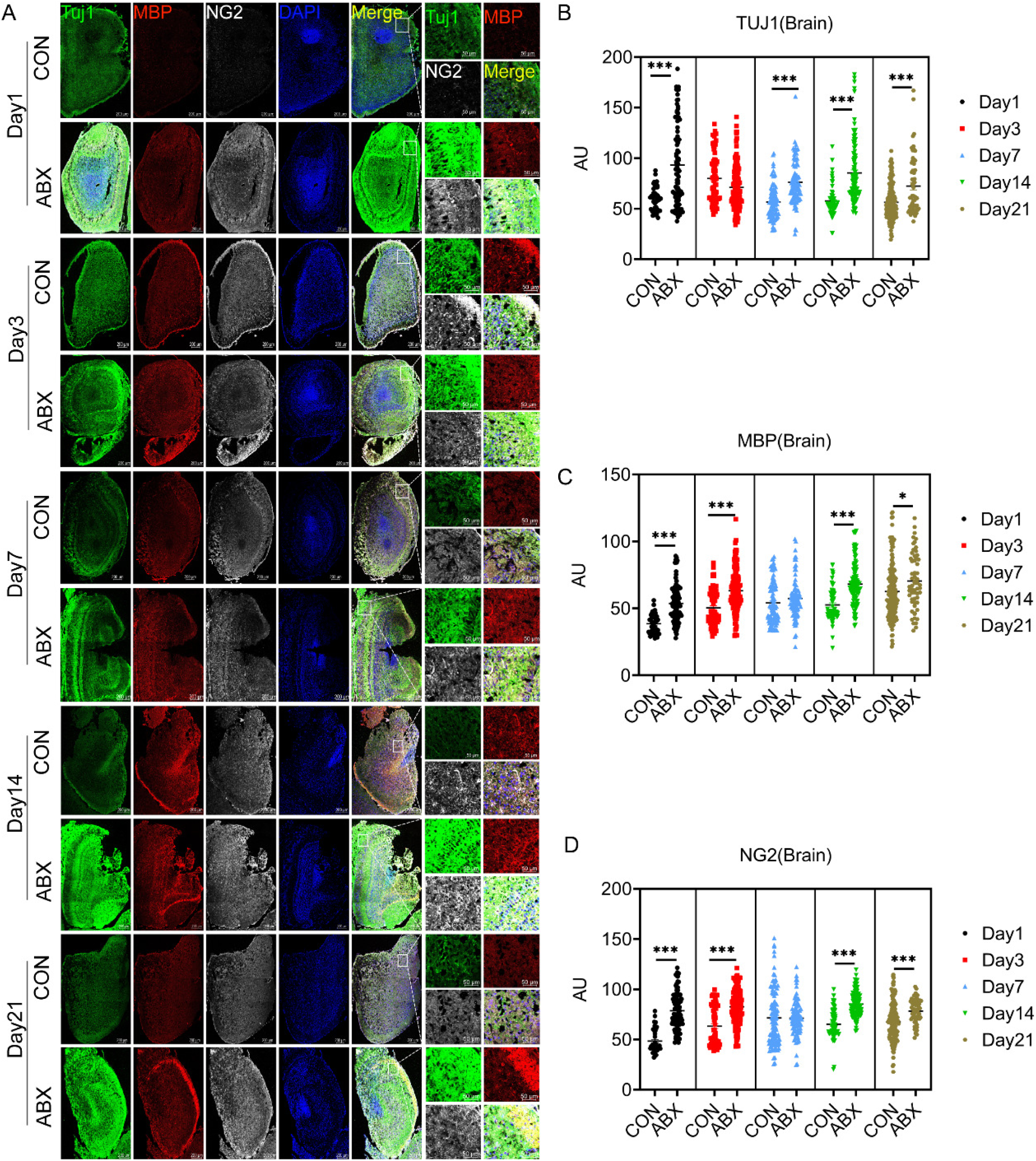
Maternal exposure to antibiotics impacted on the brain neurodevelopment a of offspring. Brain neurodevelopment in mice in CON and ABX group on the 1^st^, 3^rd^, 7^th^, 14^th^, 21^st^ days after birth (A). images of immunofluorescent staining using TUJ1 (green), MBP (red) and NG2 (white) as characteristic markers of neuron, myelin and mature oligodendrocytes, NG2 glia respectively. Scatter plots of TUJ1 (B), MBP (C) and NG2 (D) in CON and ABX group. n = 3 biological repeats. Scale bar, 200μm. *P* values for comparisons between the 2 groups in Student’s *t*-test tests. * *p*<0.05, *** *p*<0.001.

To determine whether the gut microbiota was associated with the neurodevelopment, we conducted 16S rRNA gene sequencing of fecal samples from the CON and ABX groups. For taxonomic diversity, compared with CON group, lower chao index on the 1^st^ day after birth revealed lower evenness of gut microbiota in ABX mice, while the Shannon and Simpson indexes revealed that antibiotic exposure had no significant effect on the species richness in ABX mice (**Supplementary table 1**). The presence of certain neurodevelopment associated bacterial species, such as *Bacteroidota* phylum and *Lactobacillus genus* is regarded as a health concern (30). We found that the abundance of *Bacteroidota* phylum and *Lactobacillus genus* significantly decreased in ABX group (**Figure 2A-B, Supplementary table 2**). Meanwhile, PCA and Lefse plot demonstrated the presence of distinct species in the two groups (**Figure 2C-D**). To further evaluate the general makeup of the gut microbiota across the two groups, we examined the degree of taxonomic similarity between the bacteria. The results revealed that *Proteobacteria* (57–99%) and *Firmicutes* (14–82%) were the leading phyla in CON and ABX group, respectively. Spearman correlates revealed that many of the dominant phylum and genus were negatively associated with expression of TUJ1 (**Figure 2E-F**). Especially, *Bacteroidota* phylum and *Lactobacillus genus* were negatively associated with the expression of TUJ1, NG2 and MBP (**Figure 2E-F**).

**Figure 2.**
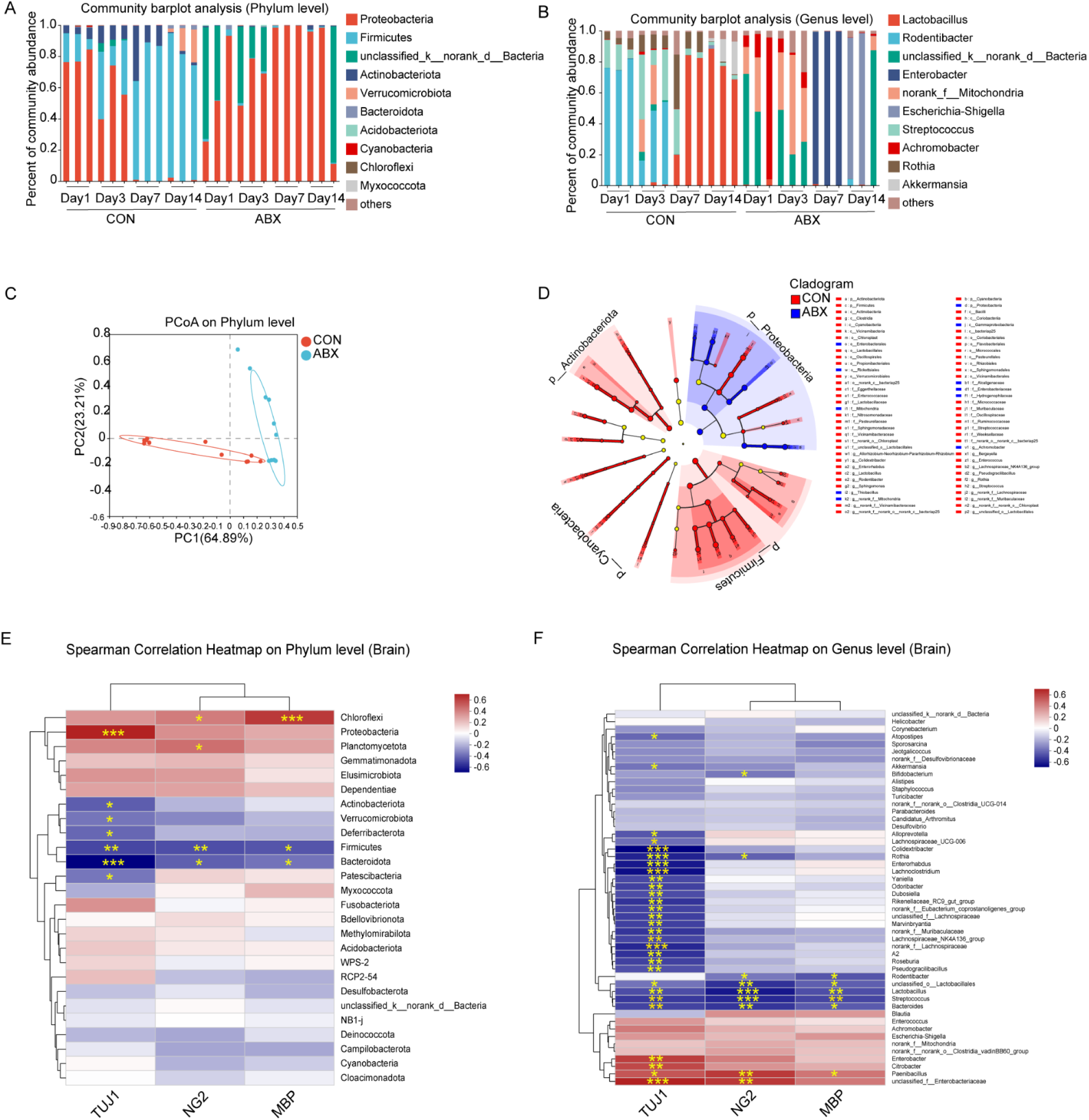
Maternal exposure to antibiotics impacted on the gut microbiota of offspring. Mice gut microbiota in phylum (A) and genus (B) level, PCoA analysis of gut microbiota (C), and Lefse cladograms (D) in control (CON) and antibiotic (ABX) group on the 1^st^, 3^rd^, 7^th^, 14^th^ days after birth. The spearman correlation heatmap between gut microbiota and neurodevelopment in brain (E, phylum level; F, genus level).

### 3.2 Maternal exposure to antibiotics impacted on the neuroregeneration after SCI and gut microbiota of offspring

Compared with sham group, sham+ABX mice had lower expression of TUJ1 (**Figure 3A-B**), and no significant change in MBP and NG2 (**Figure 3A, 3C-D**). We further investigated the role of gut microbiota in the neuroregeneration after SCI. Compared with SCI group, mice in SCI+ABX group had lower expression of TUJ1 in lesion area of spinal cord on the 1dpi, but higher on 3dpi and 14dpi (**Figure 3A-B**), along with higher expression of MBP and NG2 on the 1dpi and 3dpi (**Figure 3A, 3C-D**), and smaller injury area of the lesion core on the 1dpi (**Figure 3E**). Compared with SCI group, SCI+ABX mice had lower evenness and richness of the gut microbiota, with lower chao index on the 7dpi and 14dpi, and lower Shannon index on the 7dpi (**Supplementary table 3**). The abundance of *Bacteroidota* phylum and *Lactobacillus genus* decreased in SCI+ABX group (**Figure 4A-B, Supplementary table 4**). PCA analysis and Lefse showed distinct species in the two groups (**Figure 4C-D**). *Firmicutes* (77–91%) and *Proteobacteria* (54–99%) and were the leading phyla in SCI and SCI+ABX group, respectively. While *Bacteroidota* phylum and *Lactobacillus genus* were non-significantly associated with TUJ1, NG2 or MBP expression in spinal cord after SCI (**Figure 4E-F**).

**Figure 3.**
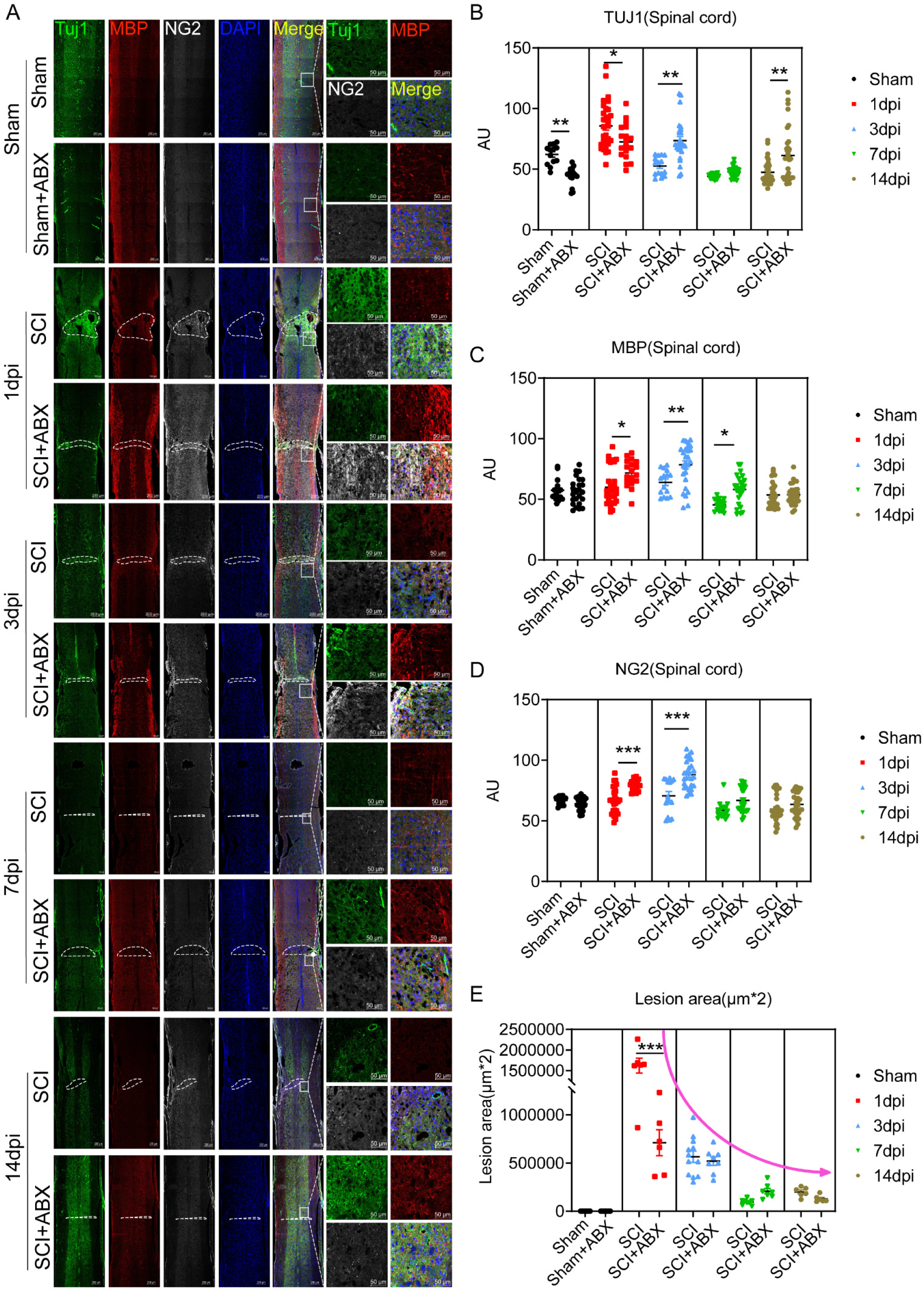
Maternal exposure to antibiotics impacted on the neuroregeneration after SCI of offspring. Neurorepair and neuroregeneration in mice in SCI and SCI+ABX group on the 1^st^, 3^rd^, 7^th^, 14^th^ days after SCI (A). images of immunofluorescent staining using TUJ1 (green), MBP (red) and NG2 (white) as characteristic markers of neuron, myelin and mature oligodendrocytes, NG2 glia respectively. Scatter plots of TUJ1 (B), MBP (C), NG2 (D), and lesion area (E) in SCI and SCI+ABX group. n = 3 biological repeats. Scale bar, 200μm. *P* values for comparisons between the 2 groups in Student’s *t*-test tests. * *p*<0.05, ** *p*<0.01, *** *p*<0.001.

**Figure 4.**
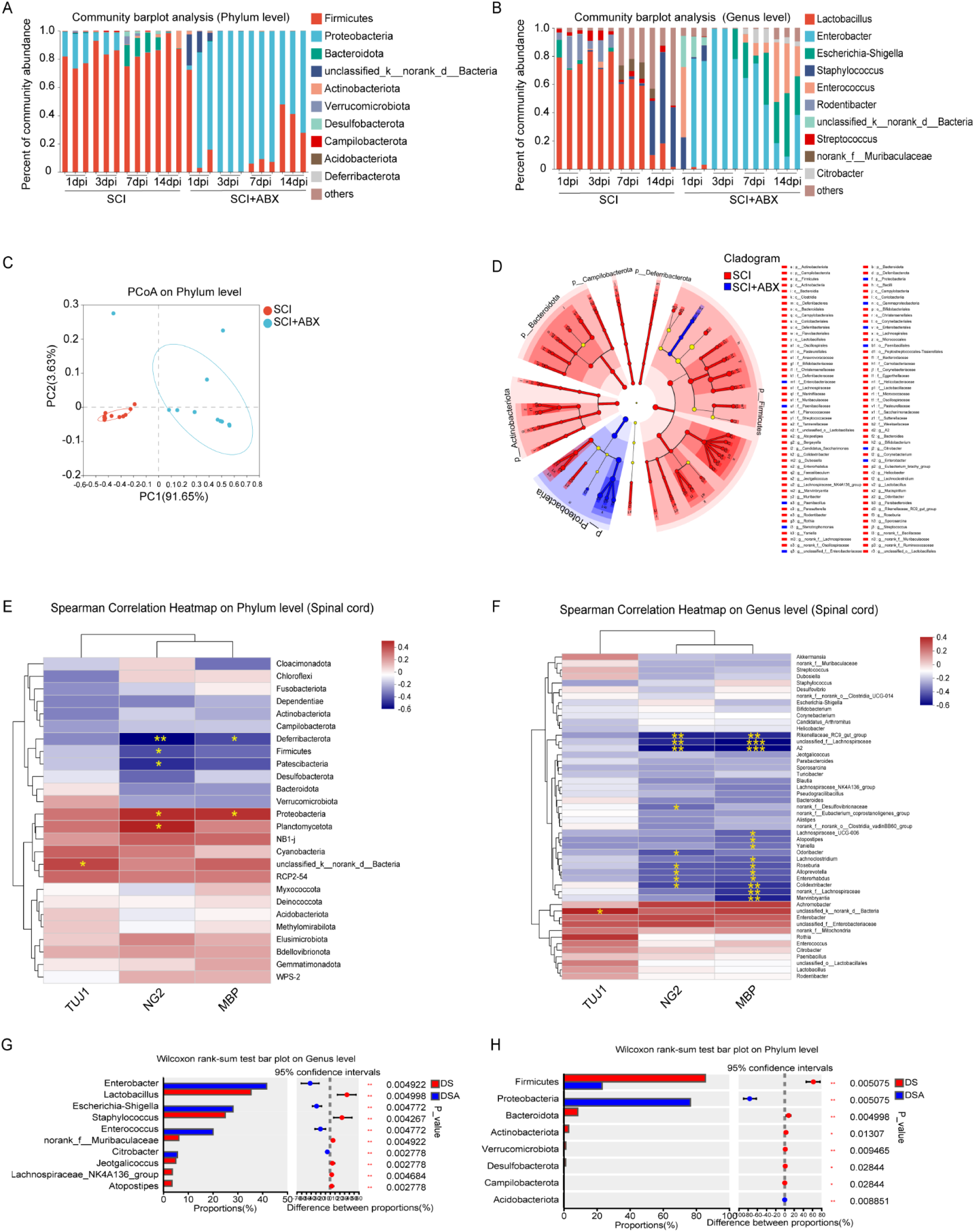
Maternal exposure to antibiotics impacted on the gut microbiota of offspring after SCI. Mice gut microbiota in phylum (A) and genus (B) level, PCoA analysis of gut microbiota (C), and Lefse cladograms (D) in SCI and SCI+ABX group on the 1^st^, 3^rd^, 7^th^, 14^th^ days after SCI. The spearman correlation heatmap between gut microbiota and neurorepair in spinal cord (C, phylum level; D, genus level)

### 3.3 Maternal exposure to antibiotics impacted on the gut metabolite composition in offspring

We found 647 differential metabolites (229 upregulated and 482 downregulated) between the SCI+ABX and SCI groups (**Figure 5A and Supplementary Figure 2-3**). Among which, Cyclo(Arg-Gly-Asp-D-Phe-Val), Ampicilloyl, DG(8:0/i-19:0/0:0), Petasinine and 6-Hydroxymelatonin glucuronide were the top 5 prominent metabolites downregulated in SCI+ABX group (**Figure 5B**). Furthermore, the KEGG enrichment analysis showed that the differential metabolites primarily enriched in amino acid metabolism pathways, including alpha-Linolenic acid metabolism, Phenylalanine, tyrosine and tryptophan biosynthesis, and Linoleic acid metabolism (**Figure 5C**). Furthermore, we selected the top 30 differential metabolites via VIP and 189 differential taxa to analyze the association between gut metabolites and microbiota, and found that 6-Hydroxymelatonin glucuronide, PC(20:3/0:0), PC(20:2/0:0), Ameltolide, DG(8:0/i-19:0/0:0), Inproquone, and Chorismate were the cores of networks (**Figure 5D**). Furthermore, we selected the same 30 metabolites to analyze the association with neuroregenaration, and found the five metabolites mentioned above were both negatively associated with TUJ1 in brain (**Figure 5E**), and MBP in spinal cord (**Figure 5F**). Altogether, these results demonstrate that maternal exposure to antibiotics impacted on the gut metabolite composition in offspring, which may be closely association with neuroregenaration.

**Figure 5.**
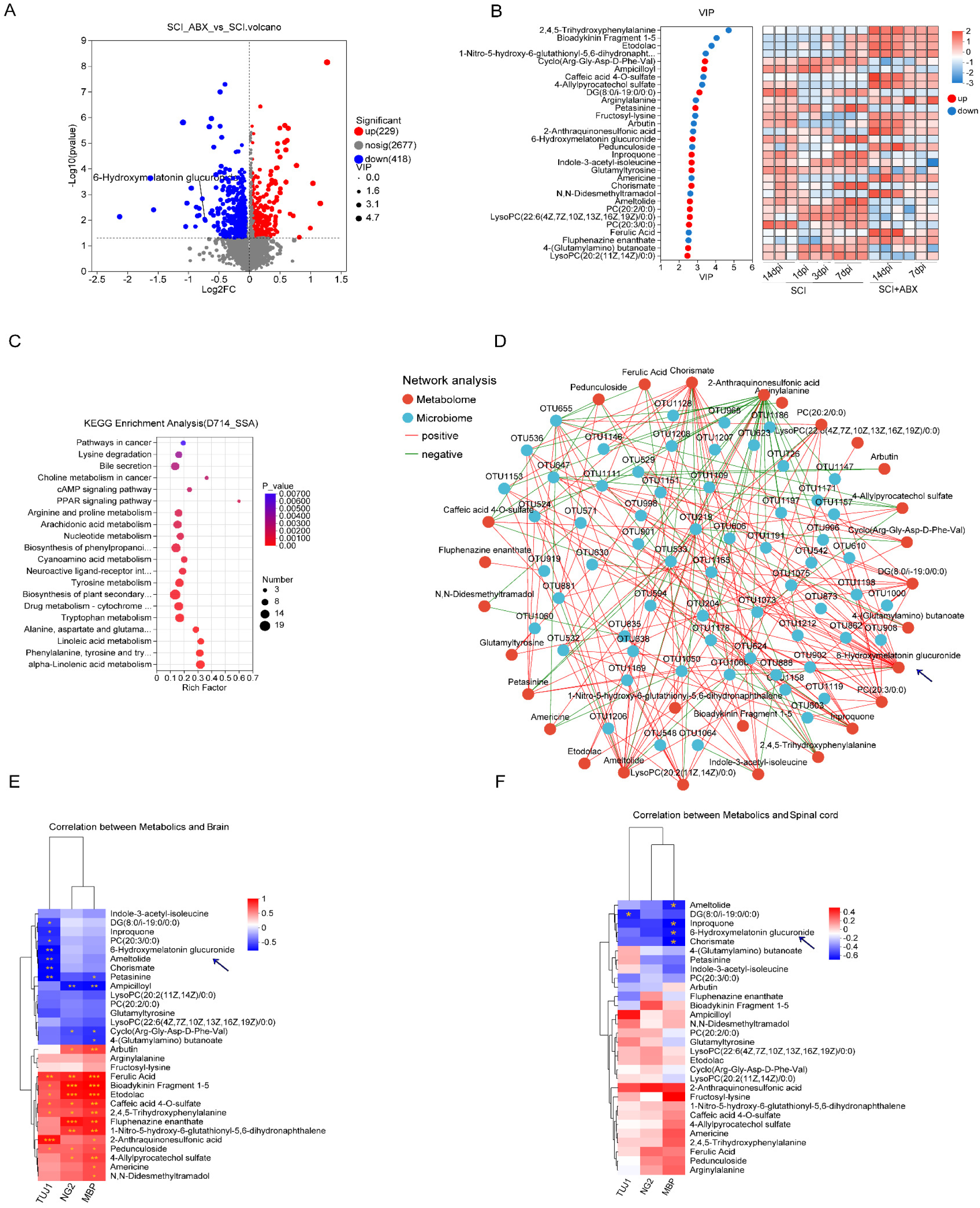
Maternal exposure to antibiotics impacted on the gut metabolite composition in offspring. (A) Volcano plot comparing different metabolites between SCI+ABX mice and SCI mice. (B) Varible importance in projection (VIP) scores of gut microbiota-derived metabolites in SCI+ABX mice and SCI mice. (C) Enriched pathway analysis of in the two groups via KEGG profile. (D) Co-expression analysis between differentially abundant microbes and metabolites. (E) Heatmap of 30 metabolites identified by VIP and markers of brain, and spinal cord (F).

### 3.4 Gut metabolites may affect NG2 glia differentiation via regulating mitochondrial fusion and fission

We found that mitochondrial fusion and fission compounds significantly altered the morphology of cultured NG2 glia cells (OLN-93) (**Figure 6A**). In particular, after the intervention of mitochondrial division inhibitor Midivi-1, the branch of OLN-93 cells was shortened and the cells were more progenitor primitive (**Figure 6A**). Meanwhile, the metabolite intervention was able to reverse this process and promoted the differentiation of NG2 glia (Figure 6A). Immunofluorescence showed that mitochondrial fusion promoter M1 and Midivi-1, reduced the expression of NG2, TUJ1, PDGFRA, MAP2 and MBP, which indicated that mitochondrial fusion and fission compounds may inhibit the differentiation of human NG2 glia (MO3.13) (**Figure 6B-E and Supplementary Figure 4**). By contrast, the metabolites intervention induced the expression of NG2, TUJ1, PDGFRA, MAP2 and MBP, indicating that the metabolite may promote the differentiation of NG2 glia (**Figure 6B-E**). Meanwhile, metabolite intervention also partially reversed the effects of M1 and Midiv-1 on the differentiation of NG2 glia (**Figure 6B-E**). These results suggested that promoting mitochondrial fusion and inhibiting mitochondrial fission may significantly inhibit the differentiation of NG2 glia, while gut microbiota metabolites could promote the differentiation of NG2 glia, partly through reversing the effects of mitochondrial fusion/fission compounds.

**Figure 6.**
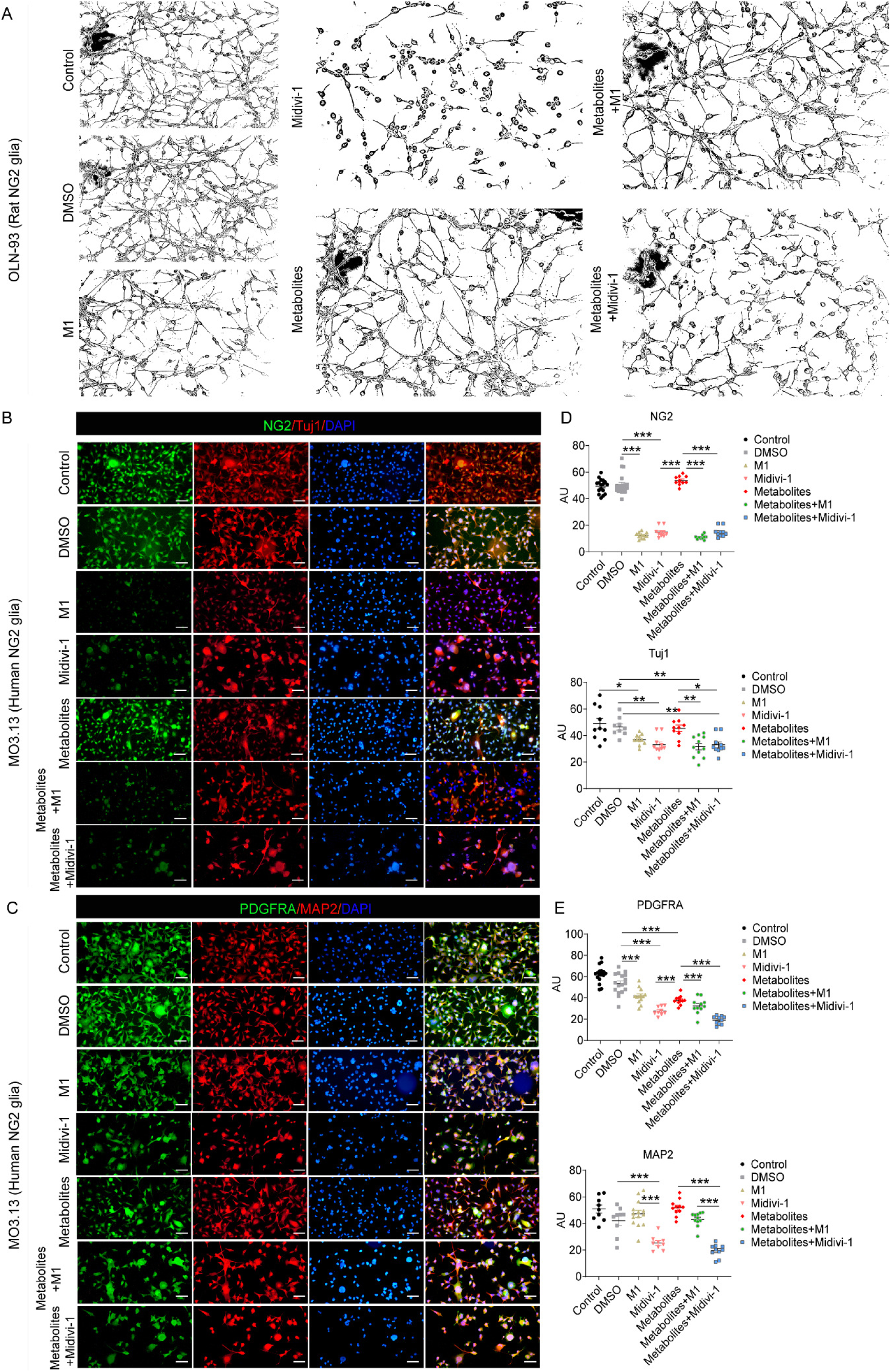
Gut metabolites may affect NG2 glia differentiation via regulating mitochondrial fusion and fission. (A) The morphology changes of cultured OLN-93 cells (Rat NG2 glia) after mitochondrial fusion and fission compounds (Midivi-1 and M1) and metabolites (TMAO+IP+HP) intervention. n = 3 biological repeats. Scale bar, 50μm. (B-C) Immunofluorescent staining of MO3.13 cells (Human, NG2 glia) by using markers of NG2 glia (NG2 and PDGFRA (green), markers of Neuron (Tuj1 and MAP2 (red)). n > 3 biological repeats. Scale bar, 50μm. (D-E), the scatter plot and statistical results of NG2, Tuj1, PDGFRA and MAP2 fluorescence intensity. *P* values for comparisons between the 2 groups in Student’s *t*-test tests. *P < 0.05, **P < 0.01, ***P < 0.001.

## 4. Discussion

In this study, we prospectively observed that antibiotic exposure (ABX) and SCI+ABX mice had higher expression of TUJ1, MBP and NG2, as well as disruption in gut microbiota and metabolites, including lower abundance of neurodevelopment-associated *Bacteroidota* phylum and *Lactobacillus genus,* and downregulated 6-Hydroxymelatonin glucuronide. Most of the key microbiota taxa and metabolites interrupted were negatively associated with TUJ1 in brain, and MBP in spinal cord. Additionally, gut microbiota metabolites promoted the differentiation of NG2 glia, partly through reversing the effects of mitochondrial fusion/fission compounds (**Figure 7**).

**Figure 7.**
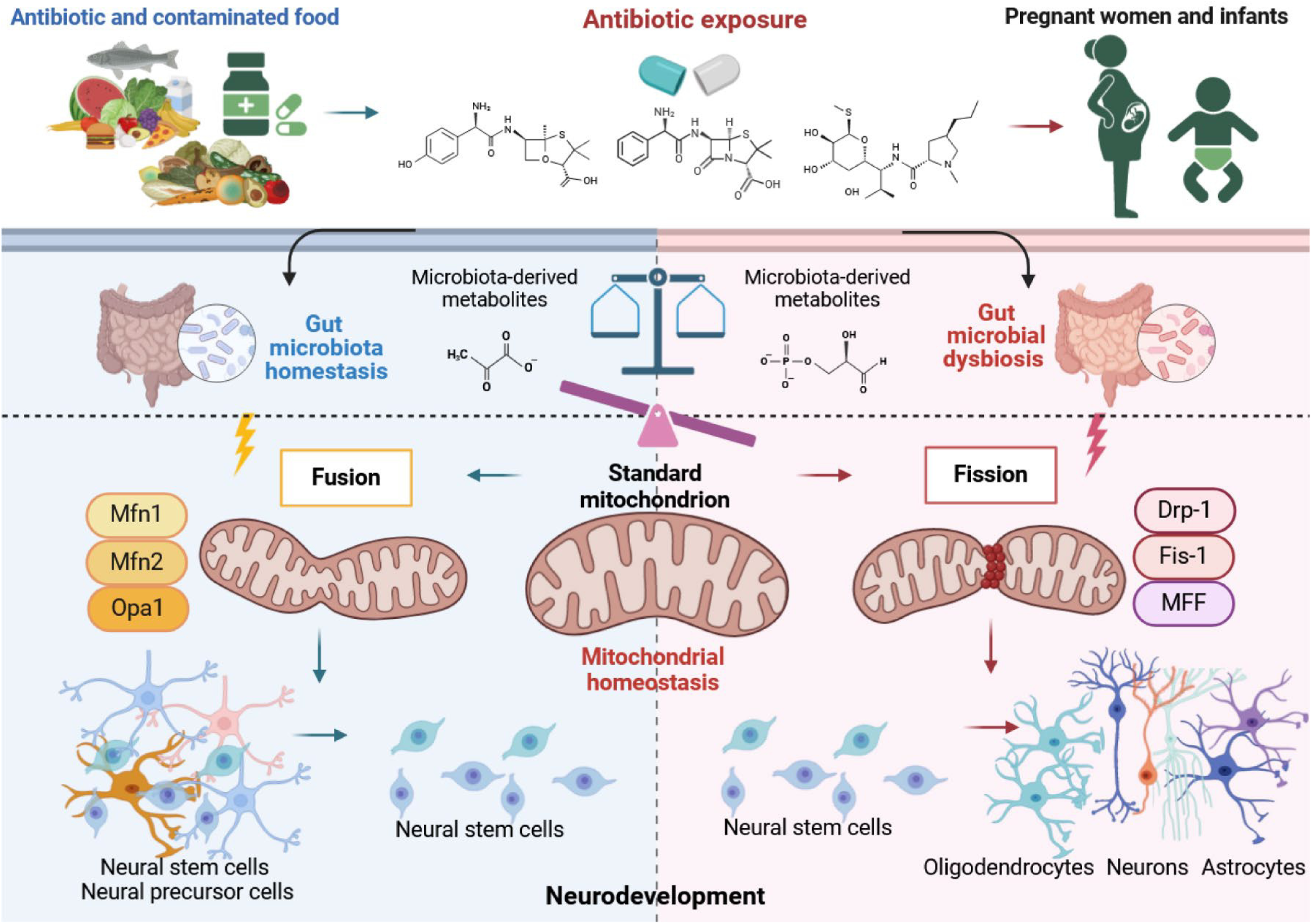
The potential schematic diagram (Created with BioRender.com). Exposure to antibiotics in early life may affect the neurodevelopment of offspring by changing the abundance of intestinal flora and subsequently mediating the metabolites of intestinal flora to interfere with mitochondrial fission/fusion homeostasis.

Our study showed that gut microbiota disruption was associated with promoted neurodevelopment in prefrontal lobe. Consistently, a previous study observed that thalamic neurons from embryos of antibiotic exposure dams increased numbers of axons, but no significant difference in axon length (11). While antibiotic exposure has been reported to decrease neurogenesis in other rodent studies (31). In a recent study, microbiota depleted mice exhibited reduced axon diameter and increased myelination in the brain (32). Brains of germ-free mice exhibited reduced expression of neuronal (NeuN), axonal (neurofilament-L), and myelination (MBP) markers relative to control mice colonized with a healthy human microbiota (33). Microbial disruption in early-life altered myelin-related gene expression in the prefrontal cortex and changed oligodendrocytes morphology in the basolateral amygdala (14). The different findings might be associated with different antibiotics intervention measures, including their concentration, type.

Findings from this work indicated that oligodendrocytes and NG2 glia, also named oligodendrocyte precursor cells (OPCs) may be involved in the interactions between microbiota with neurodevelopment and neuroregeneration after SCI. NG2 glia are with the ability to proliferate, migrate and differentiate into mature oligodendrocytes (34), which are also considered to be multipotent neural progenitor cells that could differentiate into neurons, astrocytes and oligodendrocytes (35). After SCI, oligodendrocytes and myelin are widely lost in lesion area. Successful remyelination can promote functional recovery (36) and protect axons from degeneration (37). NG2 glia are the most proliferative and dispersed population of progenitor cells in the central nervous system, and proliferate and differentiate into new oligodendrocytes to remyelinate axons after SCI (38). Consistent with our findings, one study also showed that antibiotic treatment led to the upregulation of myelin-related genes in the prefrontal cortex of neonatal mice, promoted the maturation of NG2 glia, and increased myelin production (39). Previous data support that antibiotic induced changes in the gut microbiota early after injury was associated with long-lasting oligodendrocytes activation, increased neurodegeneration of the CA3 region of the hippocampus (20). In addition, antibiotic exposure in early-life also could lead to oligodendrocytes morphology that reminiscent of the immature and malformed oligodendrocytes in the basolateral amygdala, even continued to adolescence (14). These findings signified the importance of gut-brain axis.

Omics data in our study indicated that *Lactobacillus genus,* 6-Hydroxymelatonin glucuronide and tryptophan biosynthesis pathway associated with regeneration after SCI in early-life. Melatonin, hydroxylated to 6-hydroxymelatonin and excreted in urine as the sulfate and glucuronide conjugates, is a highly neuroprotective substance that can exert cytoprotective effects (40). Except through the tryptophan-serotonin biosynthesis process in the pineal gland, the gut microbiota also contributes to the production of melatonin from tryptophan (41). Consistent with our study, a recent research observed that antibiotic-treated and germ-free mice possessed diminished melatonin levels in the serum, while fecal microbiota transplantation with *Lactobacillus reuteri* (L. R) and *Escherichia coli* (E. coli) recapitulated the effects of gut microbiota on host melatonin production by activating the TLR2/4/MyD88/NF-κB signaling pathway (42). Melatonin-treated SCI mice showed decreased relative abundance of *Clostridiales* and increased relative abundance of *Lactobacillales* and *Lactobacillus* (43). In addition, studies showed that melatonin could effectively relieve acute sleep deprivation-induced cognitive impairment (41). While the molecular mechanisms underlying remain unclear. Further studies were also needed to explore the potential effect of other metabolics found in our study.

The changes of gut microbiota in early-life coincide with the critical time course of early infant neurodevelopment. Early-life is also a critical window period for the establishment of gut microbiota, which is highly susceptible to the influence of antibiotics(10). Population studies have confirmed that the use of antibiotics can lead to long-term changes in the composition, abundance and metabolism of gut microbiota in infants and young children, which last for several years, while probiotic supplementation can reduce the effects of maternal antibiotic exposure during pregnancy on gut microbiota in infants (44). Animal studies have also found that even low-dose antibiotic exposure can cause persistent effects on the gut microbiota of mice(9). An increasing number of studies have suggested that gut microbiota can influence brain development during this critical period of early-life, leading to long-term changes in behavior (45). Microbiota metabolites are compounds produced by intestinal microbial metabolism (21). In addition to the classical hypothalamic-pituitary-epinephrine axis and endocrine pathways (i.e., intestinal peptides and hormones), metabolites produced by bacteria can affect the levels of related metabolites in the brain through blood circulation, thereby regulating brain function and development (45). The obligate anaerobic *Clostridium* (*Firmicutes*) and facultative anaerobic *Enterobacteriaceae* (*Proteobacteria*) in the gut microbiota catabolizes dietary choline to trimethylamine (TMA), which is absorbed in the gut; in turn, it is converted to trimethylamine-N-oxide (TMAO) in the liver (46). Maternal antibiotic exposure results in a significant increase in metabolites of some gut microbiota in offspring rats, especially trimethylamine oxide (TMAO), Imidazole Propionate (IP), and Hippurate (HP), which further inhibits neuronal differentiation and myelination (11).

In this study, we found that the metabolites (TMAO+IP+HP) intervention induced the expression of NG2, TUJ1, PDGFRA, MAP2 and MBP and may promote the differentiation of NG2 glia. Meanwhile, metabolite intervention also partially reversed the effects of mitochondrial fusion and fission compounds M1 and Midiv-1 on the differentiation of NG2 glia. It has been found that mitochondrial fission promotes the differentiation of Radial glia cells (RGCs) into neurons, while mitochondrial fusion maintains self-renewal of RGCs (27). Mitochondrial fusion and fission are mediated by the gtpase, and fusion is regulated by mitofusion 1 (Mfn1), mitofusion 2 (Mfn2), and optic atrophy 1 (Opa1). Fission is mainly regulated by dynamin related protein (Drp1), fission 1 (Fis1) and mitochondrial fission factor (MFF) (47). Studies have shown that Mfn2 is involved in neuronal maturation and synapse formation, and the loss of Mfn1/Mfn2 impairs the self-renewal of hippocampal neural stem cells, leading to spatial learning defects (48). The deletion of Drp1 can lead to impaired forebrain development, reduced number of neural synapses and defective synapse formation in mice (48). Although we did not the changes of mitochondrial function and genes, these results may indicate that the microbiota metabolites had partly affect the NG2 glia differentiation or neurodevelopment via crosstalk with mitochondrial fusion and fission. Unfortunately, the absent of germ-free (GF) group and without testing that whether the antibiotics treatment itself had effects on neurobehaviors of mice, we assumed that antibiotic-related neuroactivity was mediated by the gut microbiota.

## Conclusions

Antibiotic exposure in early-life leaded to significant changes in the composition, abundance, and metabolites of gut microbiota in offspring. Gut microbiota in early-life was associated with neurodevelopment in brain, and neuroregeneration after SCI, maybe partially by the synergy of *Lactobacillus genus-* tryptophan-melatonin. We also revealed that 6-Hydroxymelatonin glucuronide may be one of the core metabolites in the pathway, and gut microbiota metabolites could promote the differentiation of NG2 glia, partly through reversing the effects of mitochondrial fusion/fission compounds. Antibiotic exposure in early-life should also be obtained more attention in our daily life.

## Author’ contributions

Q.Z. and P.P.F. contributed equally to this work. Q.Z. and P.P.F. designed and performed experiments, analyzed the data and drafted the manuscript. X.Y.M, X.M.T., Y.X.Z participated in the experiments. Q.C.L., X.Z., and F.Y. conceived and designed the experiments, supervised the overall project and revised the manuscript. All authors have reviewed and approved the final manuscript as submitted and agree to be accountable for all aspects of the work.

## Availability of data and materials

Sequence data associated with this project have been deposited in the NCBI Sequence Read Archive (SRA) database (Identification number: PRJNA1039240).

---

**Supplementary Figure 1. Experimental flow chart.**

**Supplementary Figure 2. Differential metabolite analysis between SCI+ABX mice and SCI mice (7dpi).** (A) Volcano plot comparing different metabolites between SCI+ABX mice and SCI mice (7dpi). (B-C) KEGG enriched pathway analysis and the differential abundance score of KEGG pathway in the two groups. (D) Varible importance in projection (VIP) scores of gut microbiota-derived metabolites in the two groups. (D) Differential metabolite analysis between SCI+ABX mice and SCI mice. (E-F) Compounds and pathway classification of metabolite analysis between the two groups.

**Supplementary Figure 3. Differential metabolite analysis between SCI+ABX mice and SCI mice (14dpi).** (A) Volcano plot comparing different metabolites between SCI+ABX mice and SCI mice (14dpi). (B) KEGG enriched pathway analysis in the two groups. (C) Varible importance in projection (VIP) scores of gut microbiota-derived metabolites in the two groups. (D-E) Compounds and pathway classification of metabolite analysis between the two groups.

**Supplementary Figure 4. Gut metabolites may affect NG2 glia differentiation via regulating mitochondrial fusion and fission**. (A) Immunofluorescent staining of MO3.13 cells (Human, NG2 glia) by using MBP (red). n > 3 biological repeats. Scale bar, 50μm. (B), the scatter plot and statistical results of MBP fluorescence intensity. *P* values for comparisons between the 2 groups in Student’s *t*-test tests. ***P < 0.001.

